# *multideconv* - an integrative pipeline for efficiently combining first and second generation cell type deconvolution results

**DOI:** 10.1101/2025.04.29.651220

**Authors:** Marcelo Hurtado, Abdelmounim Essabbar, Leila Khajavi, Vera Pancaldi

## Abstract

The number of computational methods for cell type deconvolution from bulk RNA-seq data has been increasing in the last years, but their high feature complexity and variability of results across methods and signatures limit their utility and effectiveness for patient stratification. Applying multiple combinations of deconvolution methods and signatures often results in hundreds of redundant or contradictory cell type features describing the composition of complex tumour samples. Benchmarking efforts are inherently limited by the lack of bias-free ground truth, often yielding inconsistent results or no consensus.

To address these limitations, we present *multideconv*, an R package that reduces dimensionality and eliminates redundancy in deconvolution results, through unsupervised filtering and iterative correlation analyses. Built on top of existing frameworks, *multideconv* harmonizes outputs across methods to identify robust cell type proportion estimates and mitigate signature-driven heterogeneity. We benchmarked *multideconv* against two existing methods that provide similar functions and found it to yield more accurate estimations of cell type proportions based on virtual bulk reconstruction from single-cell expression datasets. We also increase computational efficiency by providing a meta-cell aggregation of the single-cell datasets, showing it preserves the samples’ complexity.

Despite our focus on tumour samples in the context of immuno-oncology, the tool is flexible and can be adapted to infer mixed sample composition from bulk RNAseq datasets.

The *multideconv* R package and tutorials are available at https://github.com/VeraPancaldiLab/multideconv. The code to reproduce the analysis and figures is available on github at https://github.com/VeraPancaldiLab/multideconv_paper.

## Introduction

Inferring cell type composition from complex samples, such as blood and other tissues, and its evolution across time or in disease has relevance across several biomedical fields as well as in developmental biology, but especially in immuno-oncology. The tumor microenvironment (TME), composed of immune, stromal, and other cells, plays a key role in modulating tumor progression. Despite significant progress in understanding this complex system, it remains unclear why some patients respond to specific therapies while others do not and experience recurrence. Accurately characterizing the TME is critical for understanding tumor behavior and supporting treatment decisions [1]. Current technologies, such as flow cytometry, immunohistochemistry (IHC), or single-cell RNA sequencing (scRNA-seq), provide high-resolution cellular phenotyping. However, inherent biases due to cell isolation, their high cost and technical complexity, as well as data interpretation challenges, limit their routine application in clinical settings. As a result, computational methods for estimating cell type composition from more affordable and practical bulk transcriptomics experiments have continued to gain interest in the last few years.

Reference-based deconvolution methods are commonly categorized into first- and second-generation approaches. While first-generation methods rely on static gene expression signatures derived from purified cell populations, second-generation methods leverage single-cell RNA-seq data to construct dynamic, context-specific signatures that can model a broader spectrum of cell types and states. Although more flexible, second-generation methods also introduce new challenges, including variability in single-cell references, biases in cell type annotations, inconsistent marker selection, and increased computational cost.

Recent initiatives like immunedeconv [2] and omnideconv [3, 4] have attempted to standardize and benchmark these tools, providing a single point of access to several first- and second-generation deconvolution methods for easy application to gene expression datasets. However, due to the absence of absolute bias-free ground truth, benchmarking efforts often produce discordant or context-specific outcomes, making it difficult to generalize results or define a universally “best” method. Some approaches have sought to reconcile method differences by deriving a consensus output [5], but this can oversimplify the biological complexity and fail to exploit the complementary information provided by individual tools. To capture the full complexity of the TME, it is often necessary to apply multiple combinations of methods and signatures. While this approach broadens biological insight, it also results in high-dimensional and heterogenous features that hinder interpretation and complicate downstream analysis.

To address these issues, we introduce *multideconv*, an R package designed to reduce the dimensionality and redundancy of deconvolution outputs. *multideconv* integrates the outputs of first- and second-generation methods and employs a combination of unsupervised filtering, linear correlation analysis, and proportionality-based measures to identify robust cell type features across samples. By systematically harmonizing and refining TME deconvolution features, *multideconv* enhances interpretability and facilitates patient stratification across cohorts. Built on top of existing deconvolution ecosystems and compatible with any deconvolution method, *multideconv* offers a streamlined solution for bringing coherence and interpretability in the estimate of complex tissue sample composition *(**Figure 1**)*.

**Figure 1.**
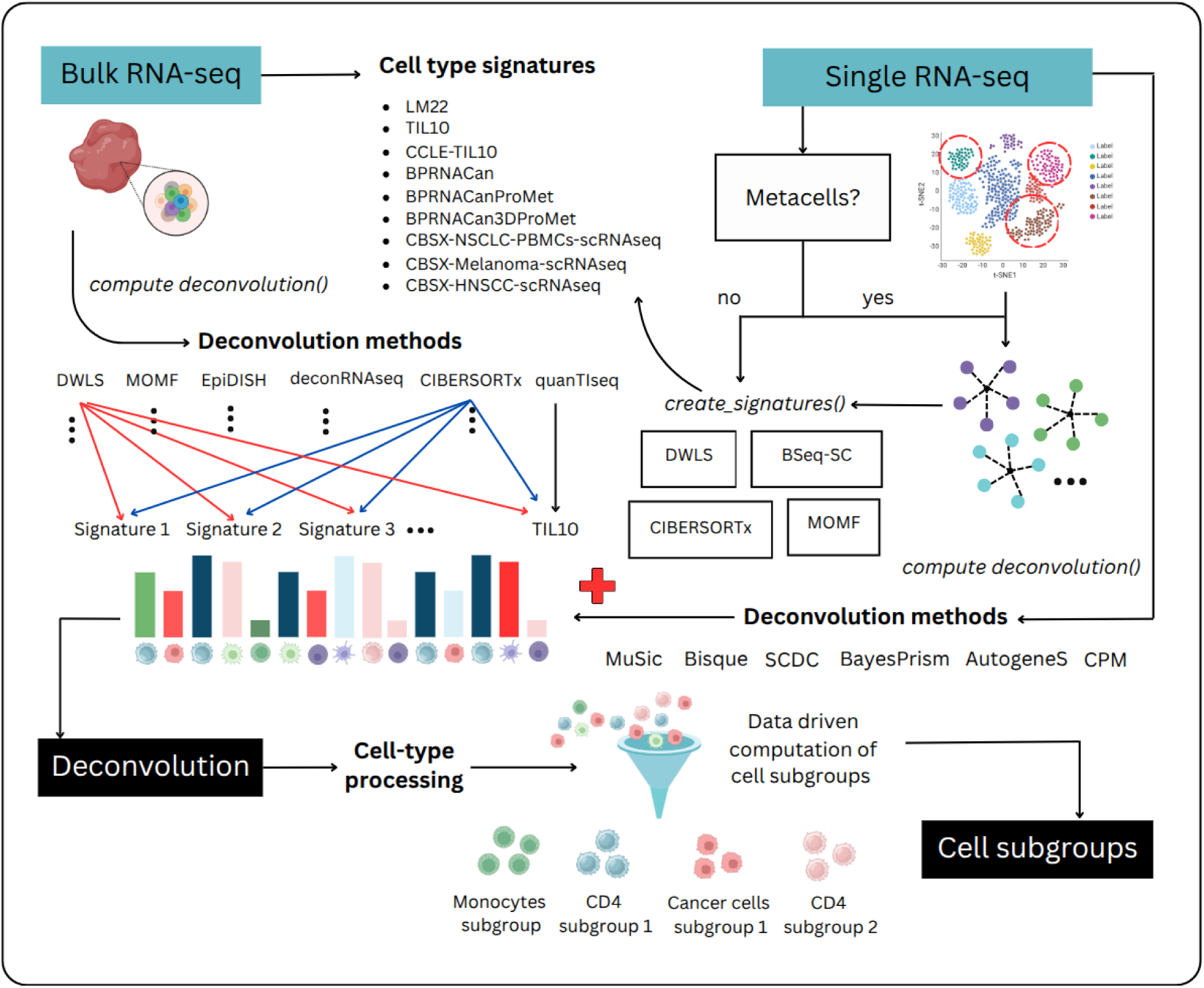
A schematic overview of the *multideconv* pipeline.

## Methods and materials

*multideconv* considers as input RNAseq raw counts with gene symbols, it performs TPM normalization followed by cell type deconvolution. By default, *multideconv* uses six deconvolution methods (CIBERSORTx, DWLS, MOMF, QuanTIseq, EpiDISH and DeconRNASeq) (***Supplementary Table 1***) combined with nine different cell signatures based on gene expression, methylation data and single cell (***Supplementary Table 2***). If scRNAseq is provided, *multideconv* also includes second-generation methods that require single cell data (MuSic, SCDC, BayesPrism, AutogeneS, CPM and Bisque) (***Supplementary Table 1***). Additionally, *multideconv* provides a function to create gene signatures using four methods (DWLS, BSeq-SC, CIBERSORTx and MOMF) (***Supplementary Table 1***) from the single cell object. These signatures will be used as additional input matrices for the six default methods.

To process the deconvolution features, *multideconv* applies a combination of unsupervised filtering techniques and iterative linear based correlations to form subgroups across all cell types (***Supplementary Figure 1***). The algorithm returns a simplified deconvolution matrix composed with subgroups of results of specific method-signature combinations based on their similarity across samples.

The first step involves unsupervised preprocessing of the deconvolution matrix. Missing values are replaced with zeros to ensure data consistency, while features with high zero occurrences (e.g., >90%) are removed, as these could bias subsequent analyses. Additionally, features with low variance across samples are filtered out, as they likely lack biological significance.

Following preprocessing, the deconvolution matrix is split into cell-type-specific groups based on established naming conventions (e.g., macrophages, NK cells, CD4/CD8 T cells) see ***Supplementary Table 3***, ensuring that features are analyzed within biologically relevant categories. Within each cell-type group, pairwise correlations are computed and highly correlated features (Spearman’s r > 0.9) are identified and pruned to reduce redundancy, retaining only one representative feature from each correlated cluster.

Subsequently, within each cell type, features are grouped into subclusters through an iterative Spearman correlation-based merging process, where pairs of features with a minimum Spearman’s r > 0.7 are progressively merged. This ensures that each subgroup contains features exhibiting strong monotonic relationships. Spearman’s correlation is chosen over Pearson’s correlation because it is more robust when different signatures may have varying numbers of cells, as it focuses on sample ranking rather than absolute values. The median value of each resulting subgroup is used as a representative feature, reducing redundancy while preserving key expression patterns. To mitigate method-specific bias, subgroups composed exclusively of features from a single deconvolution method are excluded. Each subgroup is labeled with its contributing features and methods to support downstream interpretation.

As *multideconv* includes second-generation methods that allow users to input single-cell RNA-seq data and construct cell-type signatures, these objects are often high-dimensional, sparse, and computationally demanding. To address this, *multideconv* implements a metacell construction step per cell type and patient using a k-nearest neighbors (KNN) algorithm adapted from the R package hdWGCNA [6, 7], effectively reducing the complexity of the input single-cell matrix. This task is parallelized across clusters using N workers defined by the user, significantly improving computational efficiency. Once the reduced single-cell object is built, deconvolution is performed using this compact representation, allowing for scalable integration of second-generation data while maintaining computational efficiency and performance (***Figure 1***).

## Results and discussion

We use *multideconv* to compute deconvolution of bulk RNA-seq data from bladder cancer samples [22], producing 399 deconvolution features using the default methods and signatures. After subgrouping, we are left with 114 grouped deconvolution features.

### Grouping features preserves sample clustering structure

We assessed the quality of the cell-type subgroups by comparing their silhouette scores to those derived from an unsupervised clustering approach. For each cell-type with more than ten associated features, we extracted the corresponding subset from the original deconvolution matrix and computed a correlation-based distance matrix. Hierarchical clustering was performed on this matrix, and the optimal number of clusters was determined using the gap statistic. The resulting clusters served as a baseline, and their quality was quantified using silhouette scores. We then evaluated the silhouette scores of our cell-type subgroups, computed using the same distance metric. This comparison allowed us to quantitatively assess how well our predefined clusters capture structure in the data relative to a data-driven clustering baseline. Scores are summarized in ***Table 1***, where we can see the overall silhouette scores for the subgroups are higher than the baseline, suggesting not only better cohesion, but also that grouped features may even capture more coherent biological structure than unsupervised methods.

**Table 1.**
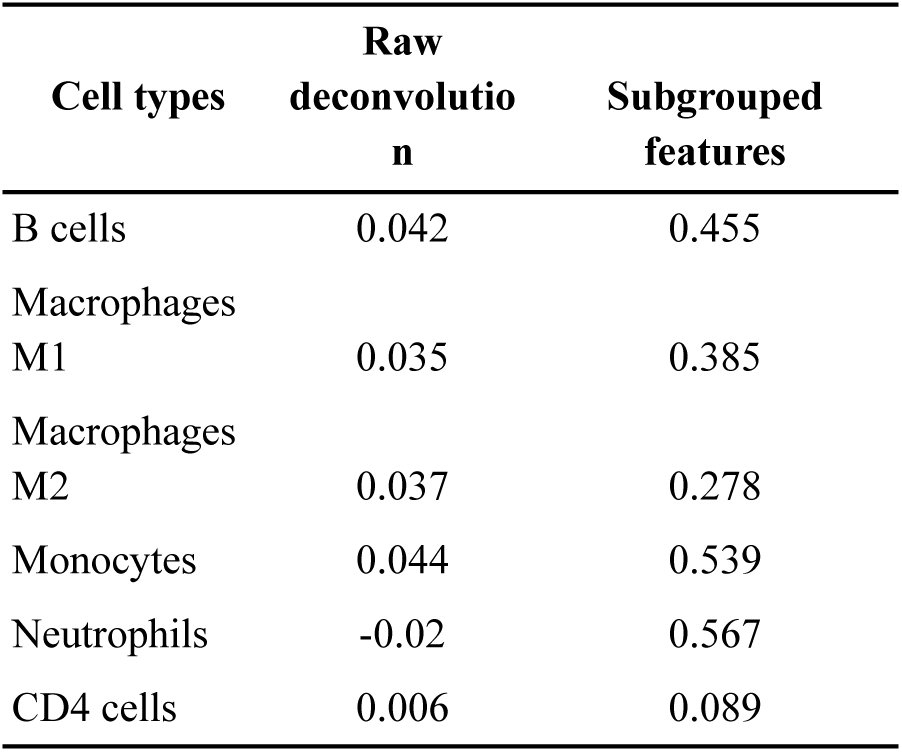
Silhouette scores.

| Cell types | Raw<br>deconvolutio<br>n | Subgrouped<br>features |
| --- | --- | --- |
| B cells | 0.042 | 0.455 |
| Macrophages<br>M1 | 0.035 | 0.385 |
| Macrophages<br>M2 | 0.037 | 0.278 |
| Monocytes | 0.044 | 0.539 |
| Neutrophils | -0.02 | 0.567 |
| CD4 cells | 0.006 | 0.089 |

### Grouping features preserves original data structure

To assess whether the algorithm preserves the underlying structure of the data after subgrouping, we performed principal component analysis (PCA) on both the original deconvolution matrix and the subgrouped feature matrix. Prior to PCA, we filtered out low-variance and low abundance features to retain informative signals. Visual comparison of PCA projections showed similar distributions between raw and subgrouped features ***(Supplementary Figure 2)***, and the variance explained by the first two principal components was comparable in both cases (raw = 20% PC1 and 11% PC2, subgrouped = 14% PC1 and 10% PC2). Additionally, k-means clustering (k = 3) applied to the top ten principal components from each dataset produced similar average silhouette scores, indicating that the subgrouping strategy maintains the structural integrity of the original data (raw = 0.0897, subgrouped = 0.1545).

### Grouped features are highly predictive of immunotherapy response and reduce computational time

Deconvolution features can be used to train machine learning models to predict clinical variables. In this line, we decided to use a bulk RNAseq dataset from a melanoma cohort [23], for a set of 73 patients with known response to immunotherapy, and assess whether there was a difference in performance between using the raw deconvolution features and the subgrouped ones. Machine learning models were trained using a repeated and stratified k-fold cross-validation (k = 5, repeats = 20), where hyperparameter tuning was performed within the cross-validation loop using grid search, optimizing the area under the ROC curve (AUROC) values. Raw model outputs were calibrated using Platt’s logistic model: out-of-fold predictions from cross-validation were used to fit a logistic regression model, which was then applied to transform the raw probabilities into calibrated scores. We benchmarked a diverse set of classification algorithms using the caret R package, spanning tree-based methods (treebag, rf, C5.0, rpart, xgbTree), linear models (glm, glmnet for LASSO and RIDGE), support vector machines (svmRadial, svmLinear), instance-based learning (knn), and linear discriminant analysis (lda).

Our results showed that overall performance is similar whether using raw deconvolution features or subgroups (***Figure 2***). In order to evaluate the feature importance we used SHAP values, which provide a measure of the contribution of each feature to the model’s predictions. For each model in the ensemble, the data was split into training and test folds using k-fold cross-validation. The model was trained on the training set, and SHAP values were computed for the test set using the kernelshap method, with the training set as the background data for the SHAP calculation. The SHAP values for each fold were aggregated by averaging across all test samples, providing a feature importance score for each variable. These values were then averaged across all cross-validation folds to obtain a robust estimate of feature importance. As observed in ***Supplementary Figure 3***, subgroups retain high SHAP values for model predictions. These results suggest that our subgrouping approach retains biological variation and produces predictive variables, while offering a more interpretable and compact representation. The machine learning model performance demonstrated that similar results can be achieved using this reduced approach with fewer features, thereby improving computational efficiency and reducing computational time.

**Figure 2.**
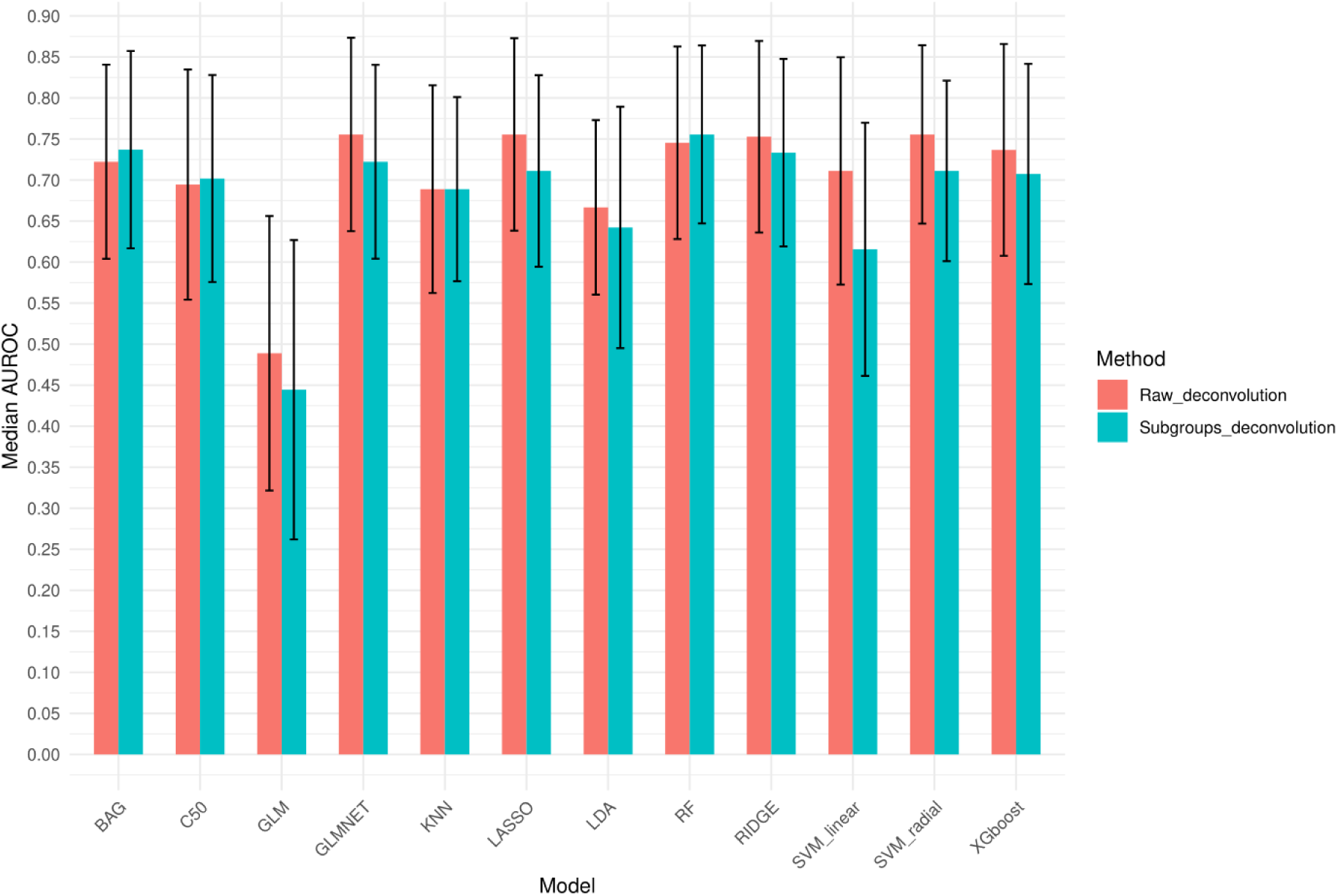
Machine learning model performance. AUROC values using raw 399 deconvolution features (red) and 114 deconvolution subgroups (blue). The bar height corresponds to the AUROC metric averaged across 5 cross-validation schemes (k = 5) with 20 repeats, and the error bar corresponds to ± 1 standard deviation, estimated across the 100 cross-validation schemes.

Notably, model training and SHAP value computation using the raw deconvolution matrix took 11 minutes and 41 minutes, respectively, compared to just 9 minutes and 4 minutes with the reduced matrix.

### Validating subgrouped features using scRNAseq datasets

To exemplify the application of *multideconv* on scRNA-seq data, we used a lung cancer dataset including 15 patients [24] ***(Supplementary Figure 4).*** We first computed metacells ***(Supplementary Figure 5)*** and used the reduced single-cell object to create static signatures before performing first- and second-generation deconvolution. To evaluate performance, we calculated pseudo-bulk profiles in the original single cell objects, essentially simulating a bulk RNAseq dataset, and deconvolved this pseudo-bulk using the specifically generated single-cell signatures and all the deconvolution methods. This procedure produced a total of 689 features and, as shown in ***Figure 3***, comparison of the deconvolution results and the real cell proportions gave an overall good result for the new signatures (DWLS r-corr = 0.95, CBSX r-corr = 0.91, MOMF r-corr = 0.9 and BSeqSC r-corr = 0.85) suggesting no loss of information during the creation and use of the reduced object for creating the signatures.

**Figure 3.**
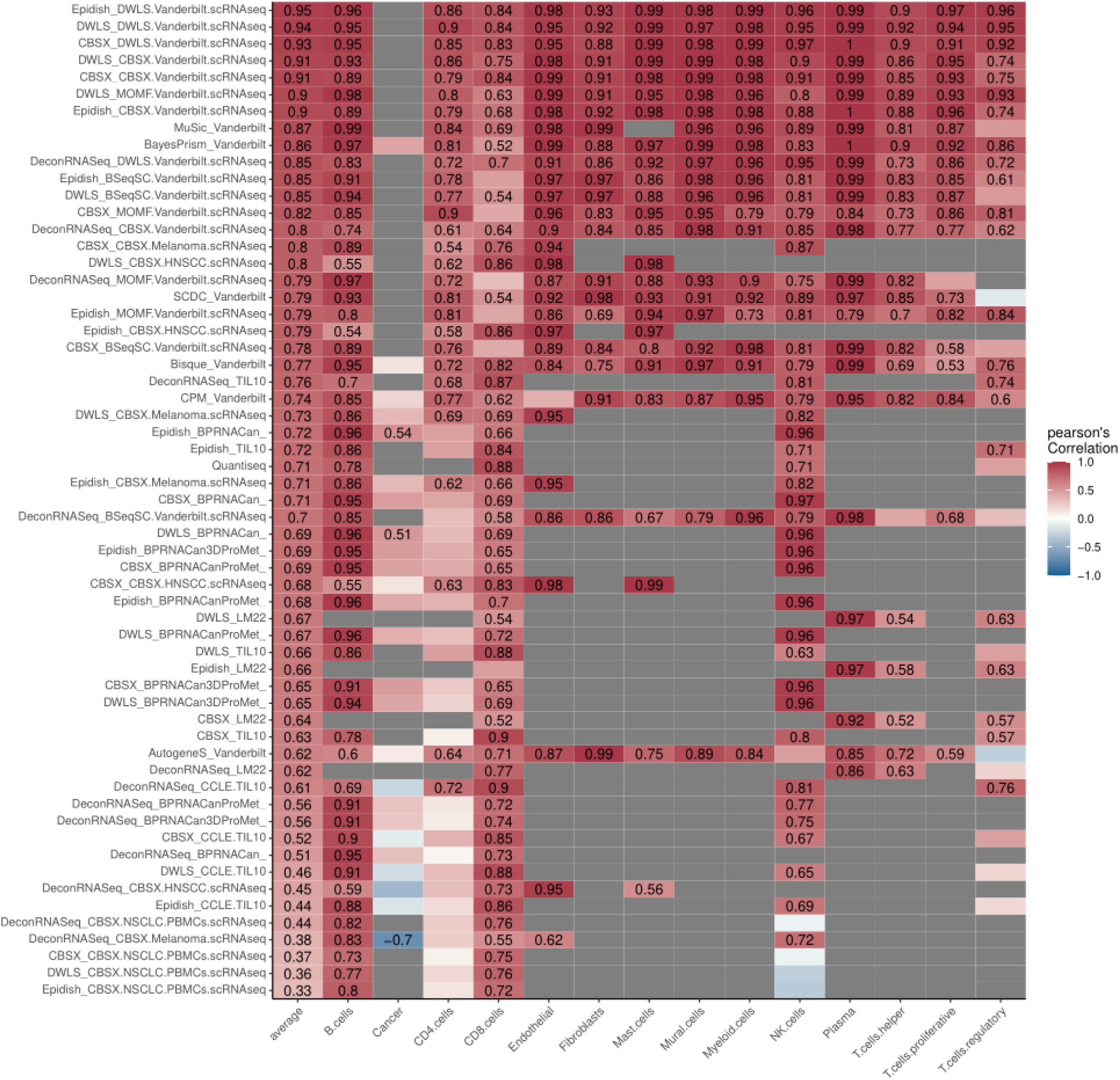
Performance of different method-signature combinations on the pseudo bulk RNAseq dataset from 15 NSCLC samples for which cell type proportion (ground truth) was estimated using the LUAD scRNAseq on the same samples [24]. Pearson’s correlations are shown as colour from −1 (blue) to +1 (red) only when p val<0.05. Non-significant correlations are left unlabeled. Grey boxes indicate cell types that were not estimated because the corresponding signature does not include that cell type. Rows indicate different deconvolution features, columns show the different cell types identified.

Additionally, we showed how the cell subgroups can produce different estimates for the same cell type from different method-signature combination, suggesting that different deconvolution results could capture potentially different cell phenotypes or states, or that they could also be affected by specific biases (***Supplementary Figure 6***).

### Benchmarking against public deconvolution aggregation methods

To evaluate the performance of our algorithm, we compared it with two publicly available methods that also perform aggregation of deconvolution results. Although these methods do not employ the same underlying approach as *multideconv*, they are designed with a similar goal of integrating deconvolution outputs. Specifically, we benchmarked decosus [25] and consensusTME [5] using the pseudobulk dataset derived from the Senosain et al. cohort, as well as an additional dataset generated from the Zilionis single-cell RNA-seq data [26] (processed into pseudobulk profiles following the same previous steps described above). We assessed the aggregated estimates using the ground truth cell proportions extracted from the corresponding single-cell datasets. To ensure a fair comparison, we excluded the previously generated signatures from the analysis. On the Senosain et al. dataset, *multideconv* achieved an average performance score of 0.76, compared to 0.69 for decosus and 0.53 for consensusTME. On the Zilionis et al. dataset, the performance scores were 0.81 for *multideconv*, 0.62 for decosus, and 0.61 for consensusTME (***Figure 4***).

**Figure 4.**
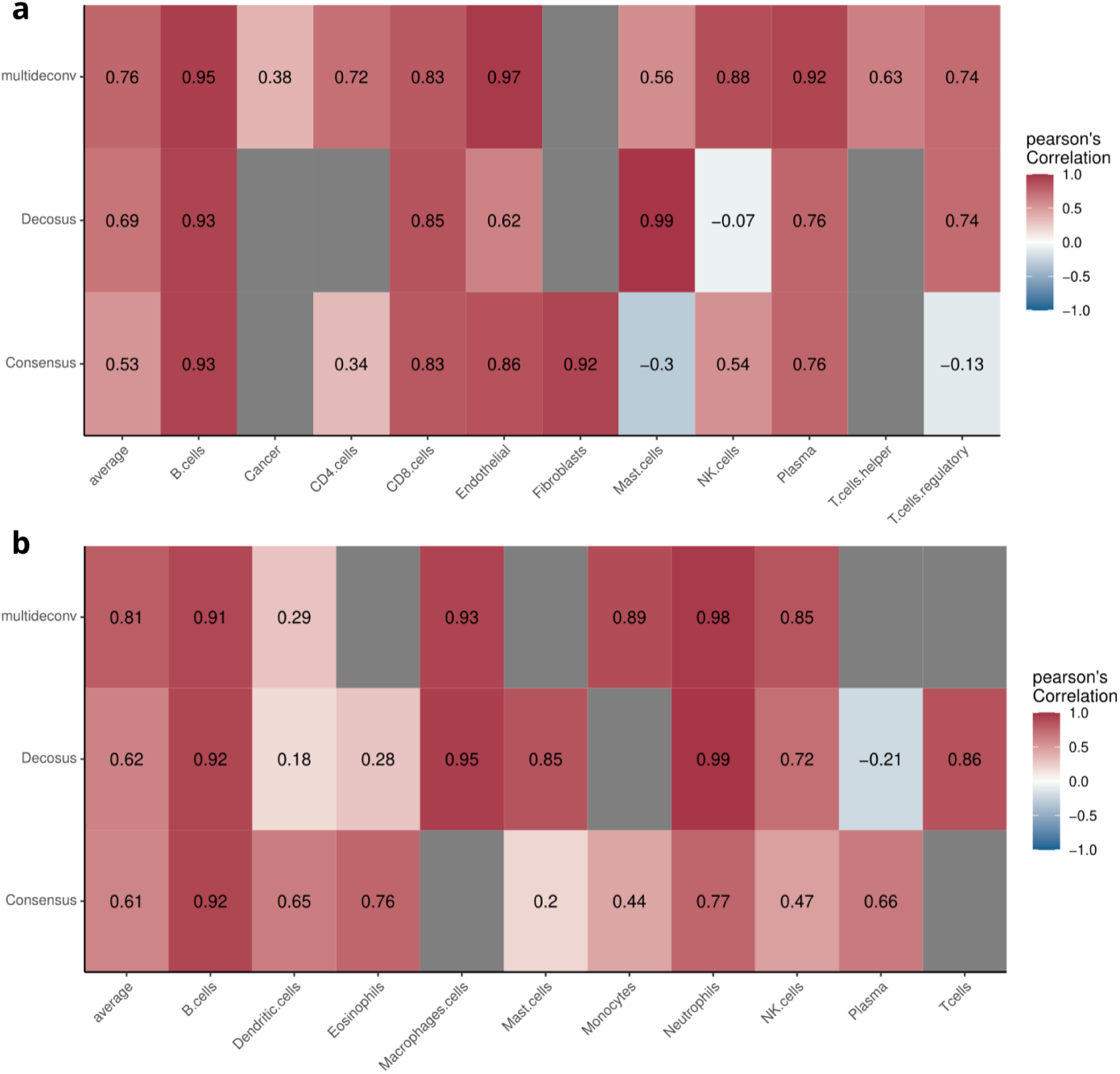
Benchmark of *multideconv* against decosus and ConsensusTME on pseudo bulk RNAseq datasets. Pearson’s correlations are shown as colour from −1 (blue) to +1 (red) only when p val<0.05. Non-significant correlations are left unlabeled. Grey boxes indicate cell types that were not estimated because the corresponding signature does not include that cell type. Rows indicate different methods, columns show the different cell types identified. **a)** Benchmark with the 15 NSCLC samples for which cell type proportion (ground truth) was estimated using the LUAD scRNAseq on the same samples [24]. **b)** Benchmark with the 7 NSCLC samples for which cell type proportion (ground truth) was estimated using the scRNAseq on the same samples [26].

## Conclusion

*multideconv* offers a practical and scalable solution for navigating the growing complexity of mixed samples deconvolution. By integrating both first- and second-generation deconvolution methods with a diverse set of static and dynamic cell-type signatures, *multideconv* transforms high-dimensional, heterogeneous features into a more biologically meaningful representation. These refined cell-type subgroups are not only easier to manage and interpret, but also better suited for downstream analyses. As the need for integrative and scalable tools continues to grow, *multideconv* offers a modular pipeline that can be easily adapted to more methods and signatures.

## Author contributions

M.H. and V.P. conceived the project. M.H. developed the pipeline and performed the analyses. L.K. and A.E. tested and provided valuable discussion on the pipeline methods. V.P. and M.H wrote and reviewed the manuscript. V.P. supervised the project.

## Acknowledgements

The authors thank Hafida Hamdache and Francesco Massaini for testing the pipeline and providing useful feedback.

## Data availability

The Mariathasan et al. datasets used to showcase the method can be found in the IMVigor210Biologies R Package. The Gide et al. bulk RNA dataset is available in the European Nucleotide Archive (ENA) under accession number PRJEB23709. The scRNAseq LUAD cohort (Vanderbilt) data is available on Zenodo under accession number 7878082. The Zilionis NSCLC scRNAseq dataset is available at GEO under the accession number GSE127465.

## Key points

- *Multideconv* reduces dimensionality and harmonizes cell type proportion estimates obtained from multiple deconvolution first- and second-generation methods and signatures, yielding interpretable immune landscapes.
- *multideconv* preserves cell deconvolution information with less features, improving clarity and interpretation.
- We demonstrate its use in stratifying patients based on cellular composition of mixed tumour samples in different cancer types
- multideconv showed superior performance compared to the few existing tools that provide similar functionality and increased computational efficiency

## Supplementary Materials

**Table S1.**
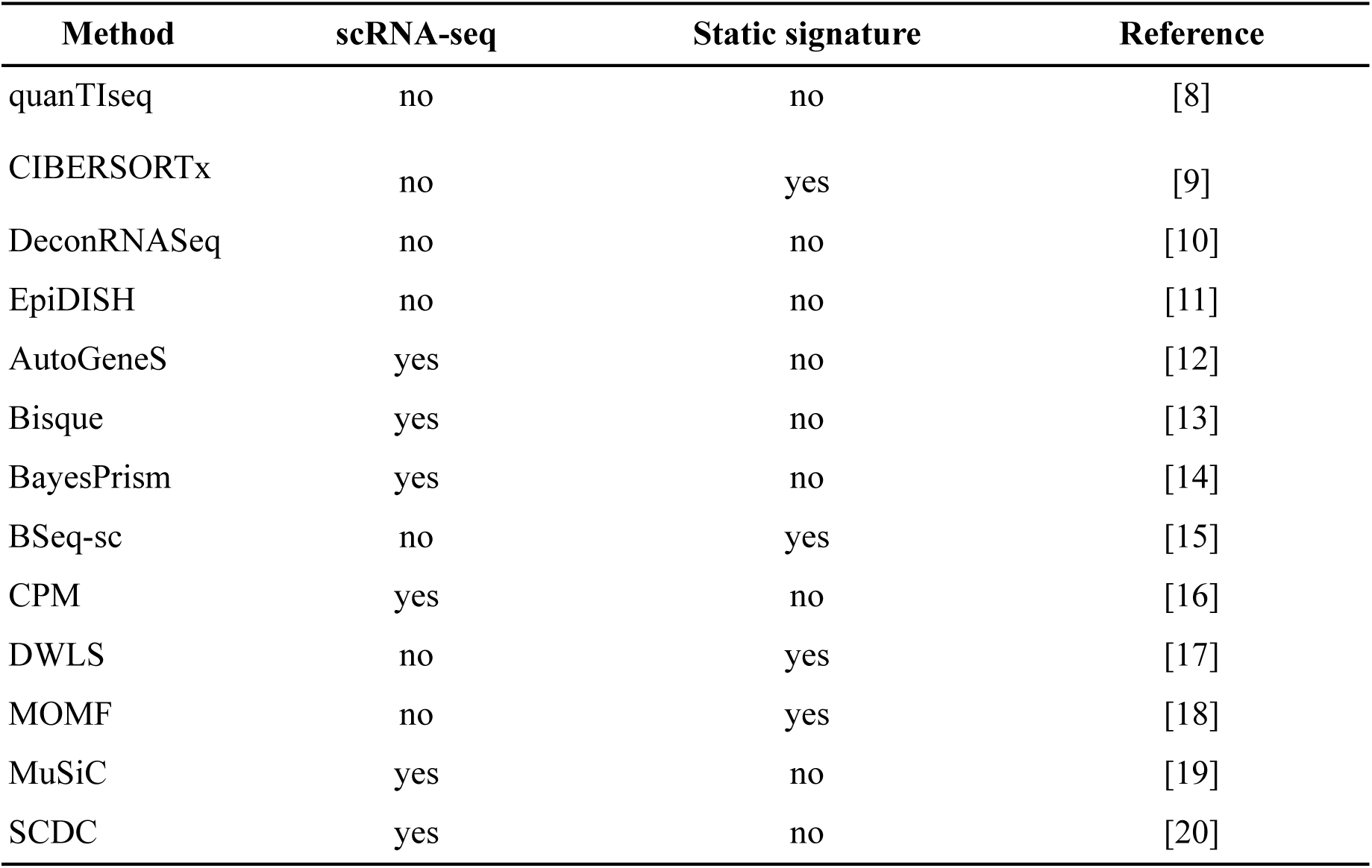
Deconvolution methods implemented in *multideconv.* Six methods need single cell data to deconvolve the bulk sample (AutoGeneS, Bisque, BayesPrism, CPM, MuSiC and SCDC) while the other seven can deconvolve the sample using external signatures with the exception of quanTIseq (CIBERSORTx, DeconRNASeq, EpiDISH, DWLS and MOMF). Four methods can be used to generate a static signature (CIBERSORTx, BSeq-sc, DWLS and MOMF).

**Table S2.**
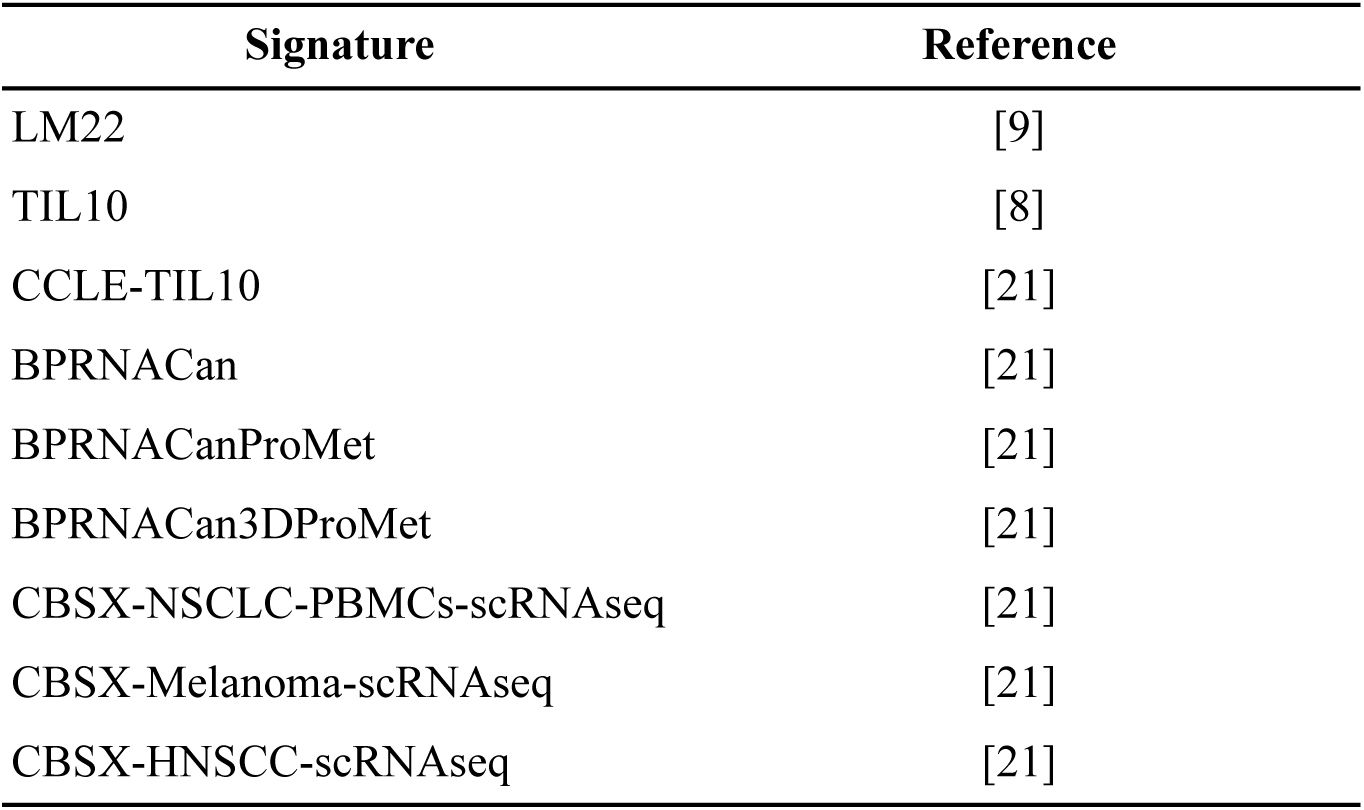
Cell type signatures provided as default in the *multideconv* package.

**Table S3.**
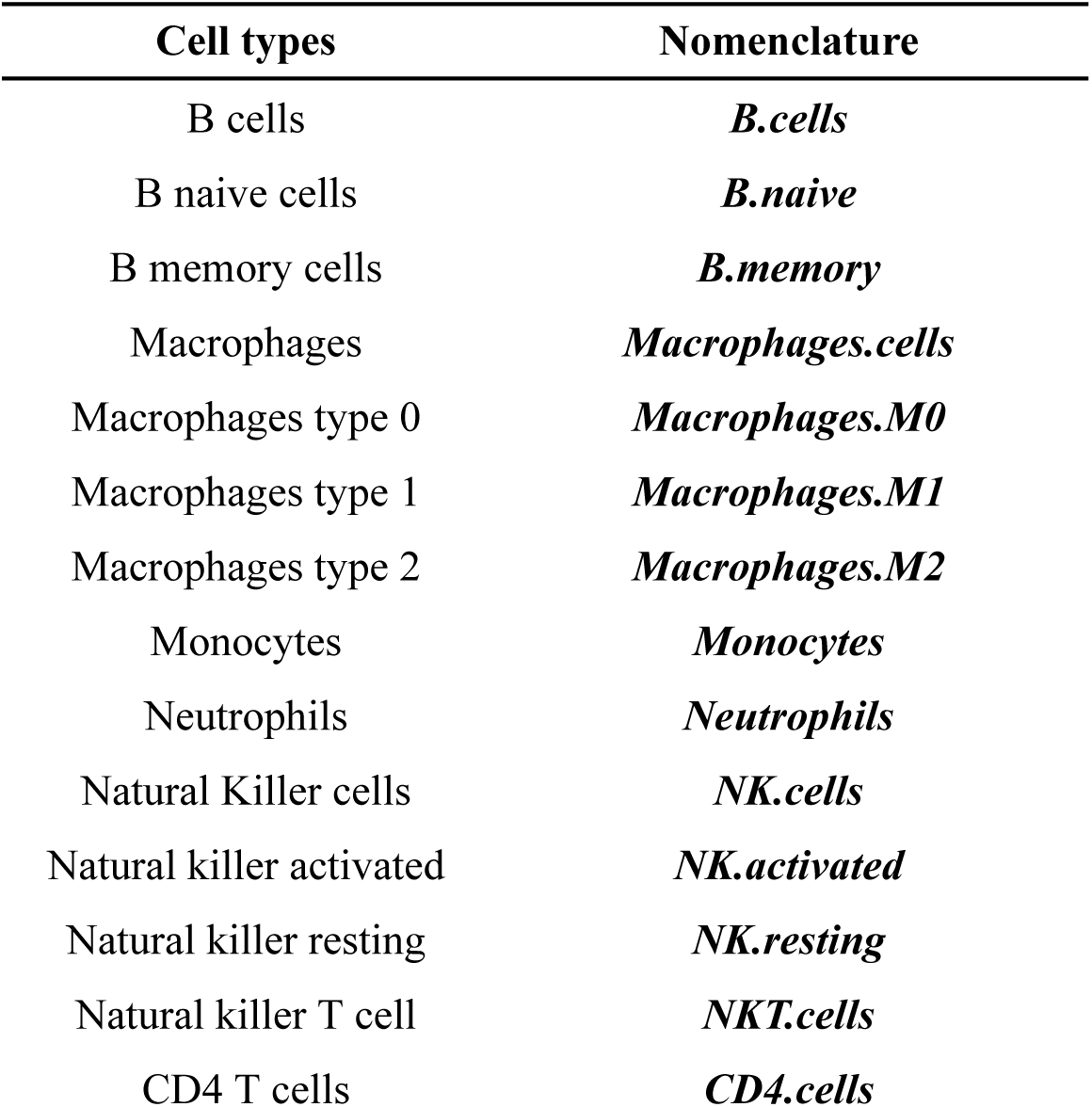
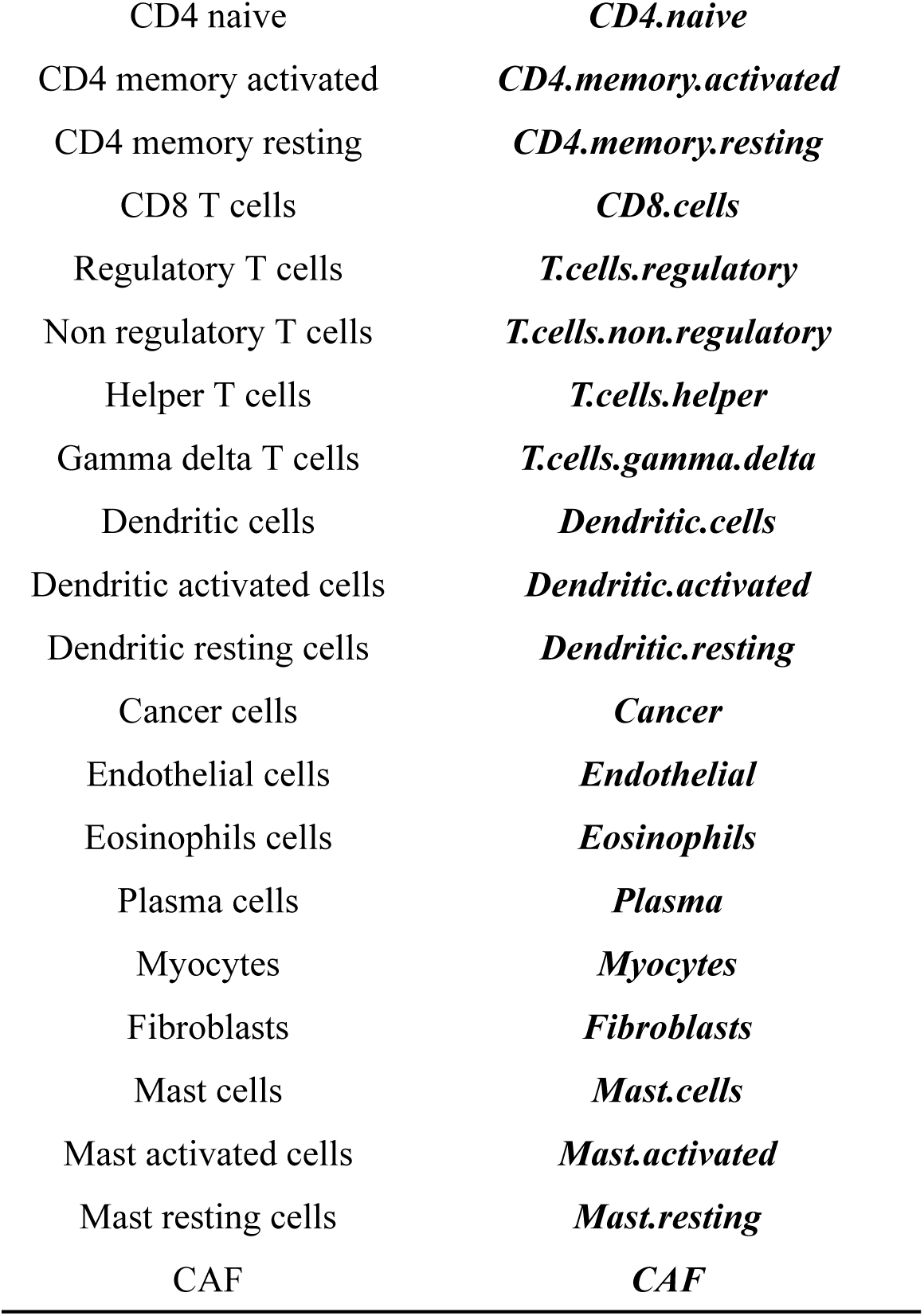
Cell types nomenclature for *multideconv*.

**Figure S1.**
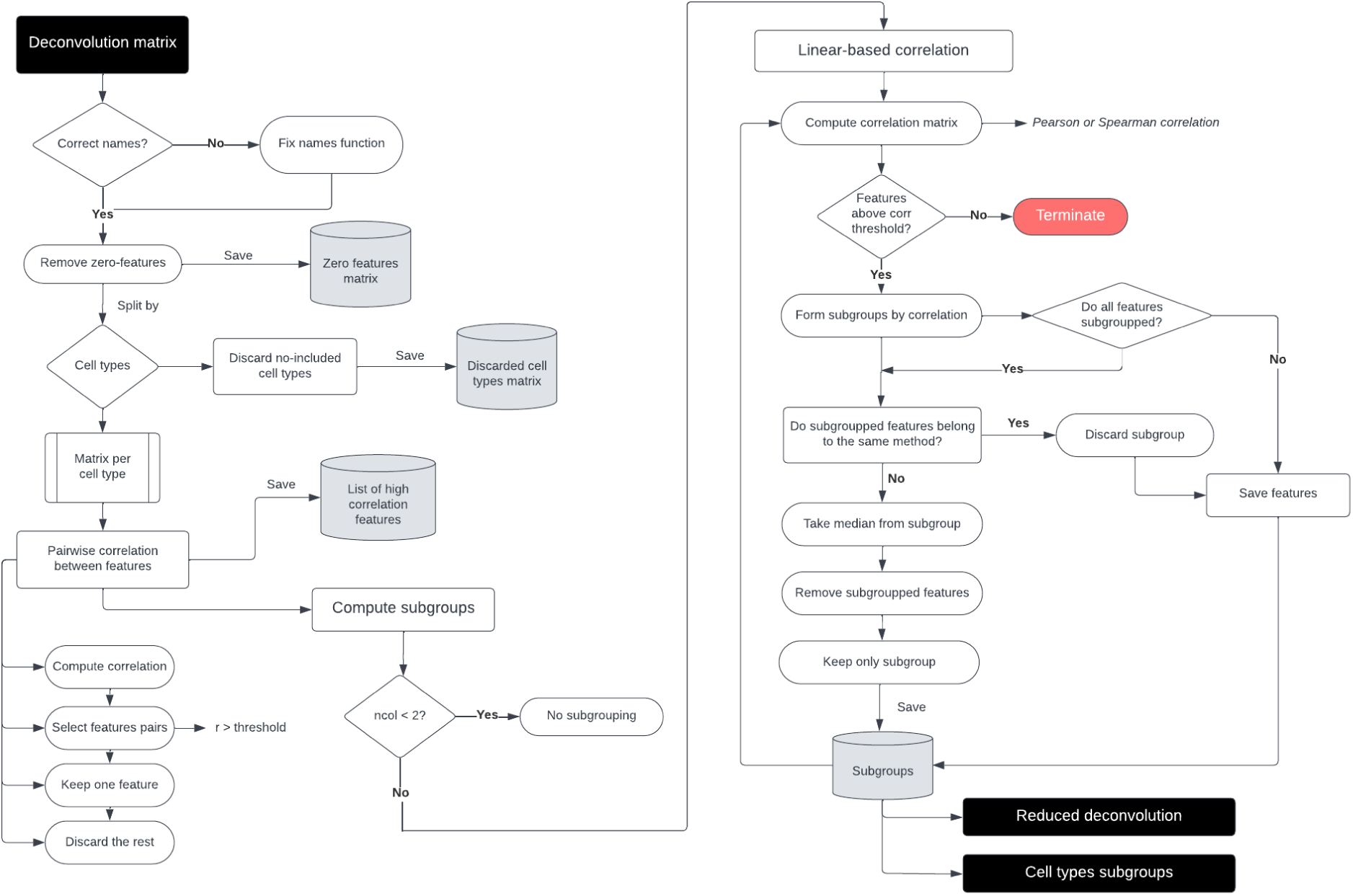
Cell-type processing algorithm.

**Figure S2.**
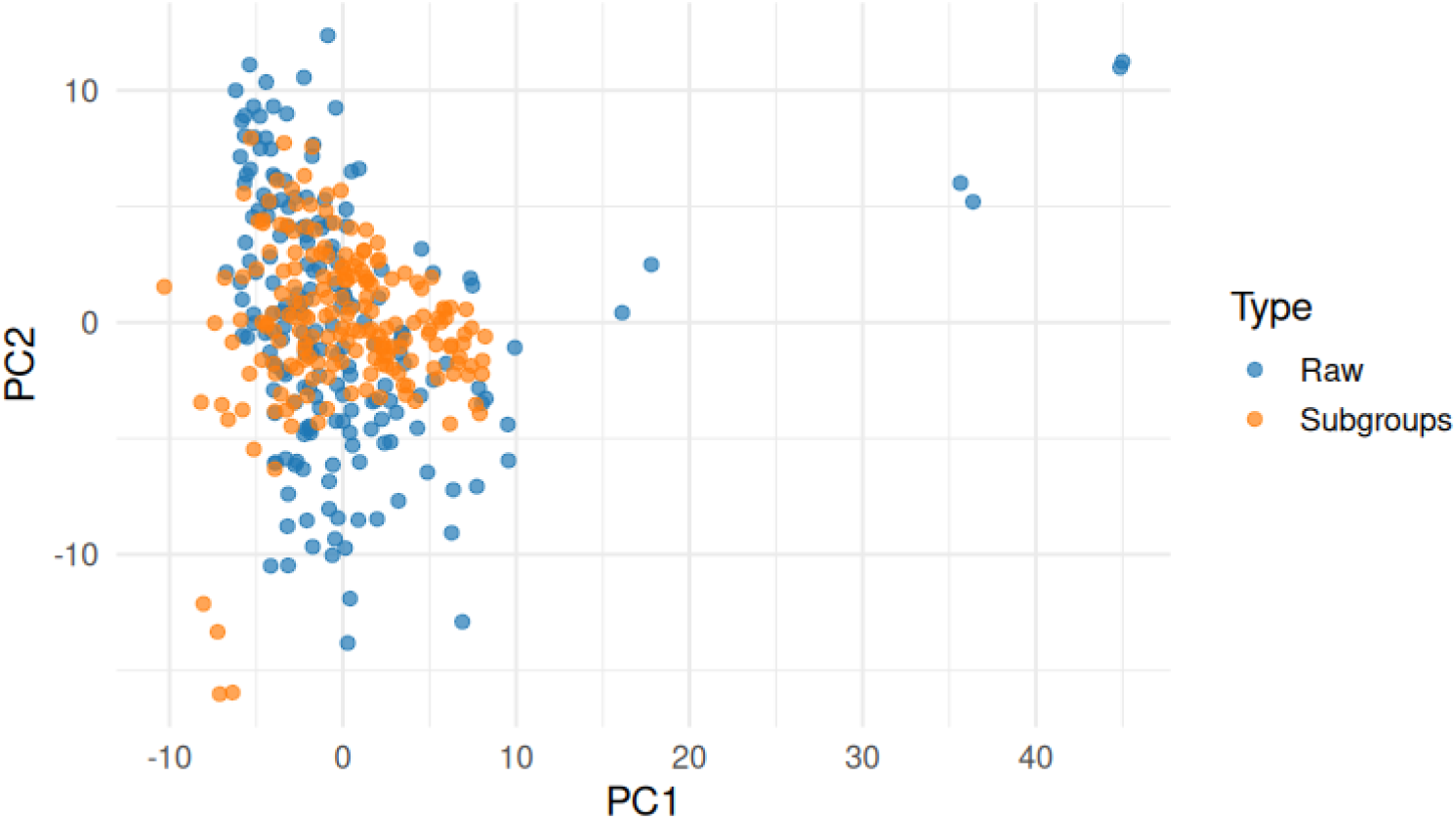
PCA using raw deconvolution matrix and subgrouped deconvolution matrix.

**Figure S3.**
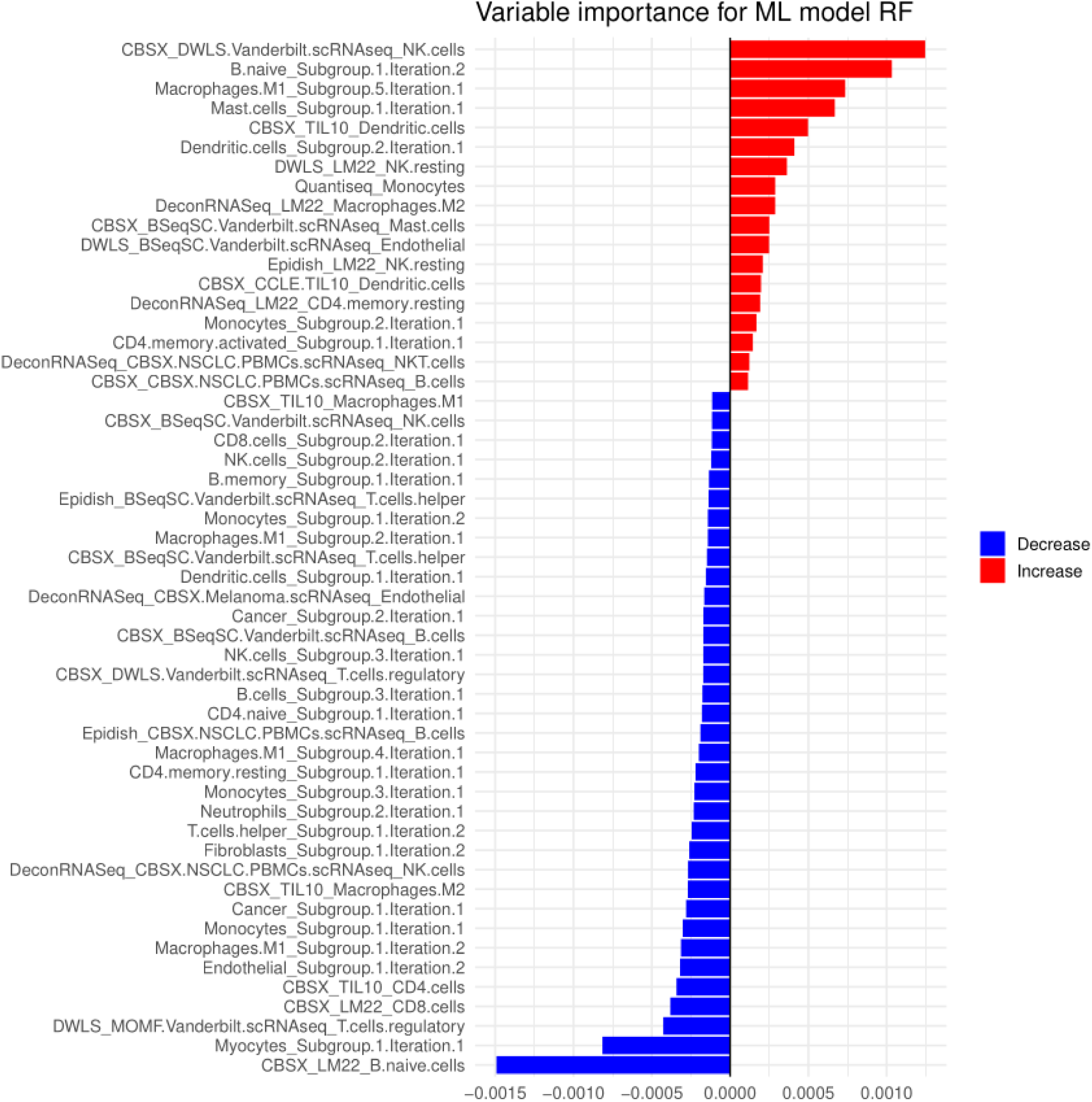
Feature importance ranking for the prediction of immunotherapy response using the subgrouped deconvolution features.

**Figure S4.**
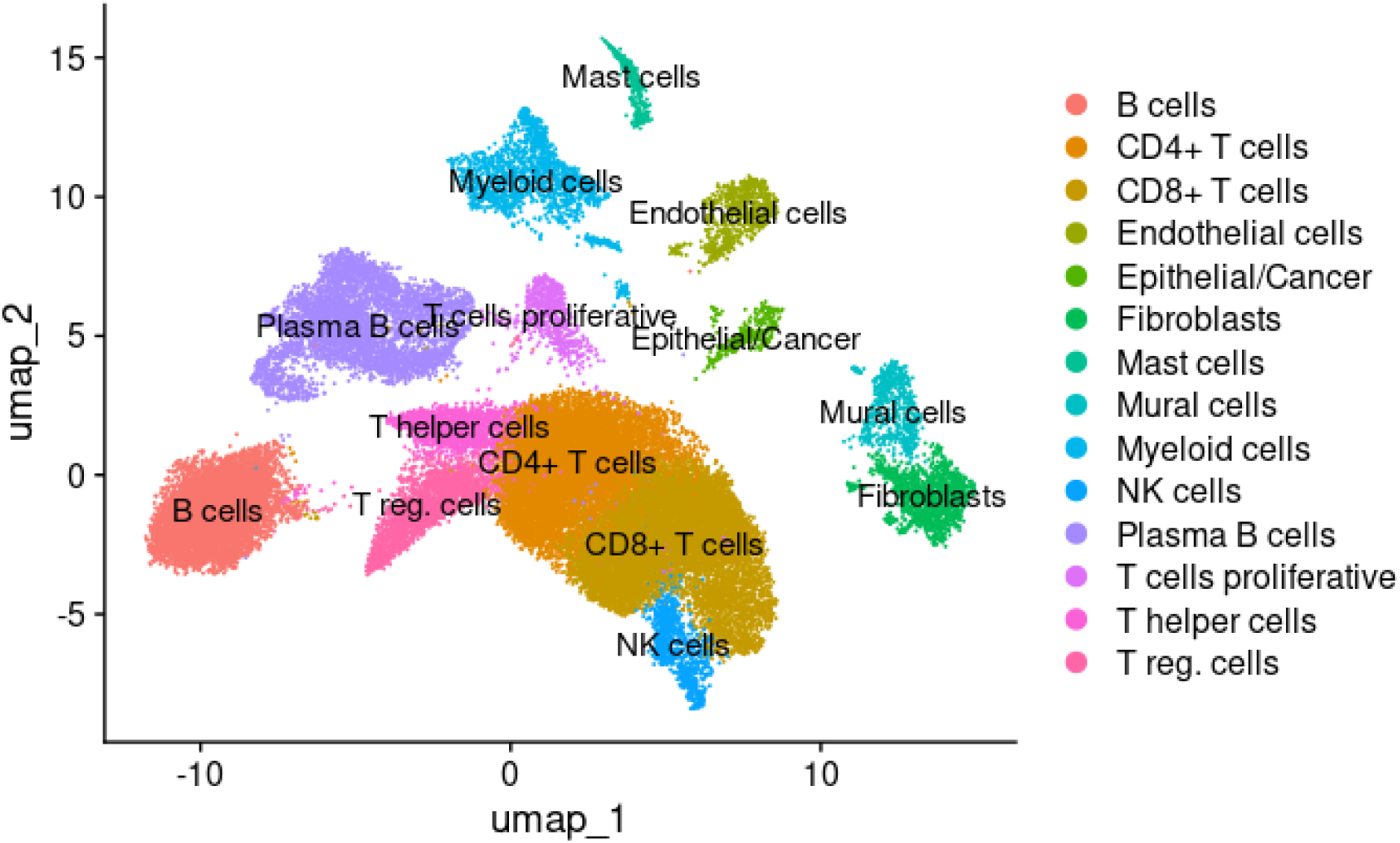
UMAP of original scRNA object from the Vanderbilt dataset [24]

**Figure S5.**
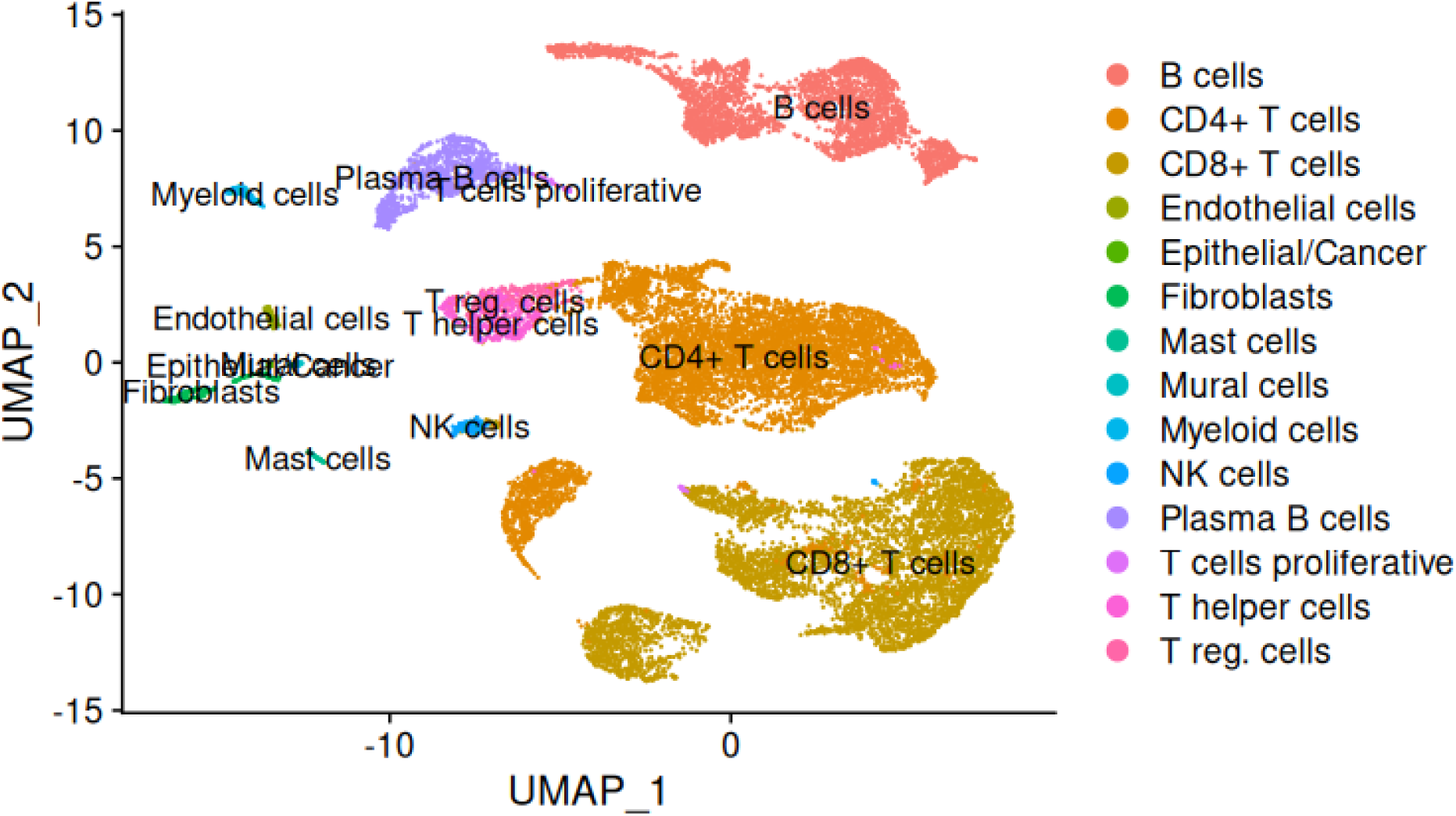
UMAP of scRNA object from the Vanderbilt dataset [24] after metacells creation

**Figure S6.**
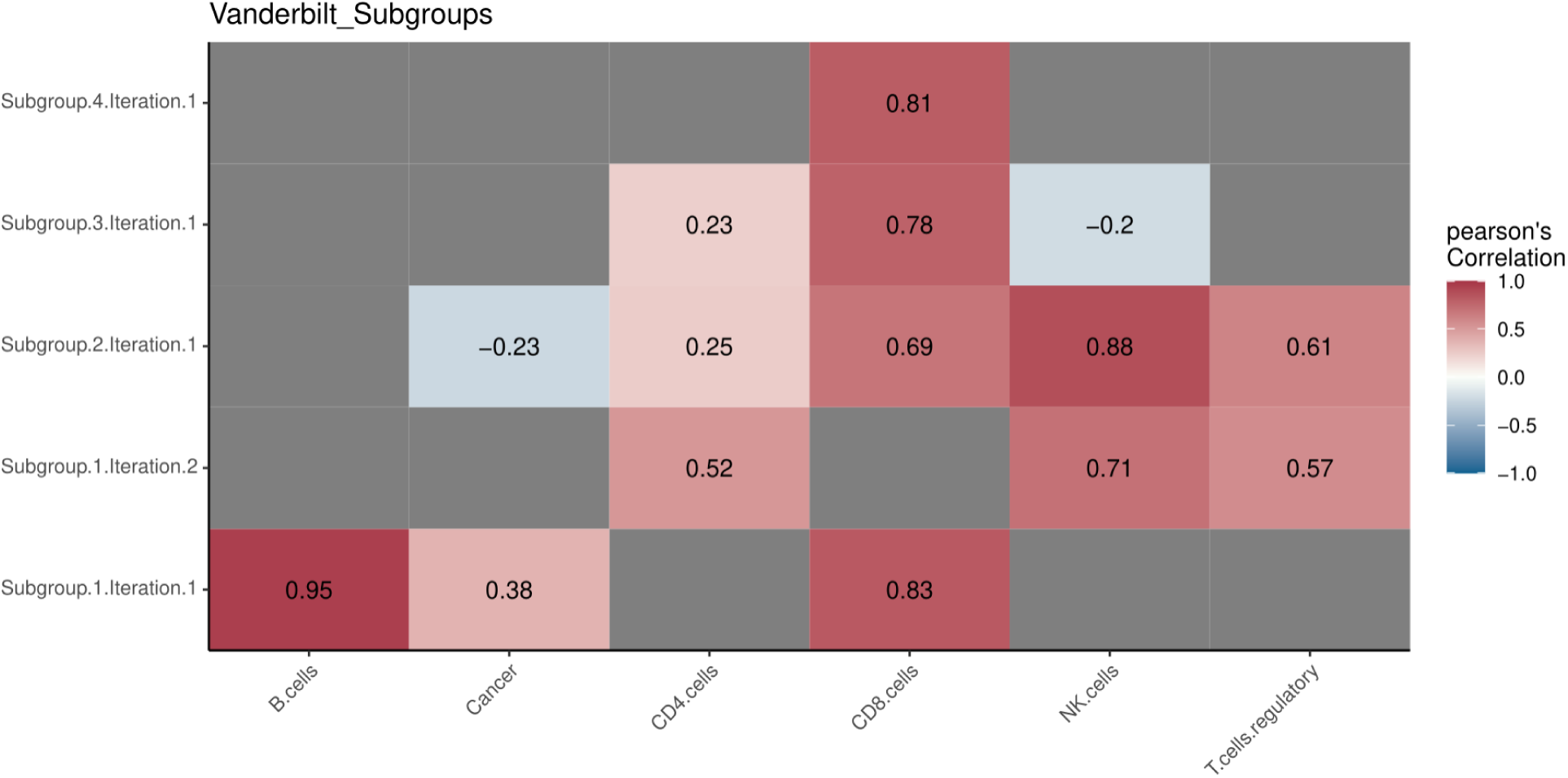
Performance of different cell subgroups estimated by *multideconv* (Y-axis) against real cell proportions estimates from the scRNA object of the Vanderbilt dataset on the X-axis [24]. Pearson’s correlations are shown as colour from −1 (blue) to +1 (red) only when p val<0.05. Non-significant correlations are left unlabeled. Grey boxes indicate cell types that were not estimated because the corresponding signature does not include that cell type.

## References

[1] Hurtado M et al. (2024). Transcriptomics profiling of the non-small cell lung cancer microenvironment across disease stages reveals dual immune cell-type behaviors. Front. Immunol. 15:1394965.

[2] Sturm, G., Finotello, F., Petitprez, F. et al. (2019). Comprehensive evaluation of transcriptome-based cell-type quantification methods for immuno-oncology. Bioinformatics, 35(14), i436–i445.

[3] Dietrich, A., Merotto, L., Pelz, K., et al. (2024) Benchmarking second-generation methods for cell-type deconvolution of transcriptomic data.

[4] Vathrakokoili, A., Zhichao, M., Yilimaz, O. et al. (2024) CATD: a reproducible pipeline for selecting cell-type deconvolution methods across tissues, Bioinformatics Advances, Volume 4, Issue 1, vbae048.

[5] Jiménez-Sánchez, A., Cast, O., & Miller, M. L. (2019). Comprehensive Benchmarking and Integration of Tumor Microenvironment Cell Estimation Methods. Cancer Research, 79(24), 6238–6246.

[6] Morabito S, Reese F, Rahimzadeh N et al. (2023). hdWGCNA identifies co-expression networks in high-dimensional transcriptomics data. Cell Rep Methods. 2023 Jun 12;3(6):100498.

[7] Langfelder, P., Horvath, S. (2008). WGCNA: an R package for weighted correlation network analysis. BMC Bioinformatics 9, 559.

[8] Finotello, F., Mayer, C., Plattner, C. et al. (2019). Molecular and pharmacological modulators of the tumor immune contexture revealed by deconvolution of RNA-seq data. Genome medicine, 11(1), 34.

[9] Newman, A. M., Liu, C. L., Green, M. R. et al (2015). Robust enumeration of cell subsets from tissue expression profiles. Nature Methods, 12(5), 453–457.

[10] Gong T, Szustakowski JD. (2013). DeconRNASeq: a statistical framework for deconvolution of heterogeneous tissue samples based on mRNA-Seq data. Bioinformatics. 2013 Apr 15;29(8):1083–5.

[11] Zheng SC, Breeze CE, Beck S, et al (2018). “Identification of differentially methylated cell-types in Epigenome-Wide Association Studies.” Nature Methods, 15(12), 1059.

[12] Aliee, H., & Theis, F. (2021). AutoGeneS: Automatic gene selection using multi-objective optimization for RNA-seq deconvolution. 10.1101/2020.02.21.940650

[13] Jew, B., Alvarez, M., Rahmani, E., et al. (2020). Publisher Correction: Accurate estimation of cell composition in bulk expression through robust integration of single-cell information. Nature Communications, 11(1), 2891.

[14] Chu, T., Wang, Z., Pe’er, D. et al. Cell type and gene expression deconvolution with BayesPrism enables Bayesian integrative analysis across bulk and single-cell RNA sequencing in oncology. Nat Cancer 3, 505–517 (2022).

[15] Baron, M., Veres, A., Wolock, S. L. et al. (2016). A Single-Cell Transcriptomic Map of the Human and Mouse Pancreas Reveals Inter- and Intra-cell Population Structure. In Cell Systems (Vol. 3, Issue 4, pp. 346–360.e4).

[16] Frishberg, A., Peshes-Yaloz, N., Cohn, O. et al. (2019). Cell composition analysis of bulk genomics using single-cell data. Nature Methods, 16(4), 327–332.

[17] Tsoucas, D., Dong, R., Chen, H. et al. (2019). Accurate estimation of cell-type composition from gene expression data. Nature Communications, 10(1), 2975.

[18] Xifang Sun, Shiquan Sun, and Sheng Yang. An efficient and flexible method for deconvoluting bulk RNAseq data with single-cell RNAseq data, 2019, DOI: 10.5281/zenodo.3373980

[19] Wang, X., Park, J., Susztak, K. et al. (2019). Bulk tissue cell type deconvolution with multi-subject single-cell expression reference. Nature Communications, 10(1), 380.

[20] Dong, M., Thennavan, A., Urrutia, E. et al. (2020). SCDC: bulk gene expression deconvolution by multiple single-cell RNA sequencing references. Briefings in Bioinformatics.

[21] Xie, T., Solórzano, J., Madrid-Mencía, M. et al. (2023). GEM-DeCan : Improved tumor immune microenvironment profiling through novel gene expression and DNA methylation signatures predicts immunotherapy response (p. 2021.04.09.439207). bioRxiv. 10.1101/2021.04.09.439207

[22] Mariathasan, S., Turley, S.J., Nickles, D. et al. (2018). TGFβ attenuates tumour response to PD-L1 blockade by contributing to exclusion of T cells. Nature 554, 544–548.

[23] Gide T.N., Quek C., Menzies A.M. et al. Distinct Immune Cell Populations Define Response to Anti-PD-1 Monotherapy and Anti-PD-1/Anti-CTLA-4 Combined Therapy. Cancer Cell. 2019;35:238–255.

[24] Senosain MF, Zou Y., Patel K. et al. (2023). Integrated Multi-omics Analysis of Early Lung Adenocarcinoma Links Tumor Biological Features with Predicted Indolence or Aggressiveness. Cancer Res Commun. 2023 Jul 26;3(7):1350–1365.

[25] Anene CA, Taggart E, Harwood CA et al. Decosus: An R Framework for Universal Integration of Cell Proportion Estimation Methods. Front Genet. 2022 Apr 1;13:802838.

[26] Zilionis R., Engblom C., Pfirschke C. et al. (2019). Single-Cell Transcriptomics of Human and Mouse Lung Cancers Reveals Conserved Myeloid Populations across Individuals and Species. Immunity. 2019 May 21;50(5):1317–1334.e10.

