## Supplementary Material for "*multideconv* - an integrative pipeline for efficiently combining first and second generation cell type deconvolution results"

### *Supplementary Materials*

#### ***multideconv* - Integrative pipeline for cell type deconvolution from bulk RNAseq using first and second generation methods**

**Table S1.** Deconvolution methods implemented in *multideconv.* Six methods need single cell data to deconvolve the bulk sample (AutoGeneS, Bisque, BayesPrism, CPM, MuSiC and SCDC) while the other seven can deconvolve the sample using external signatures with the exception of quanTIseq (CIBERSORTx, DeconRNASeq, EpiDISH, DWLS and MOMF). Four methods can be used to generate a static signature (CIBERSORTx, BSeq-sc, DWLS and MOMF).

| Method | scRNA-seq | Static signature | Reference |
|---|---|---|---|
| quanTIseq | no | no | [8] |
| CIBERSORTx | no | yes | [9] |
| DeconRNASeq | no | no | [10] |
| EpiDISH | no | no | [11] |
| AutoGeneS | yes | no | [12] |
| Bisque | yes | no | [13] |
| BayesPrism | yes | no | [14] |
| BSeq-sc | no | yes | [15] |
| CPM | yes | no | [16] |
| DWLS | no | yes | [17] |
| MOMF | no | yes | [18] |
| MuSiC | yes | no | [19] |
| SCDC | yes | no | [20] |

**Table S2.** Cell type signatures provided as default in the *multideconv* package.

| Signature | Reference |
|---|---|
| LM22 | [9] |
| TIL10 | [8] |
| CCLE-TIL10 | [21] |
| BPRNACan | [21] |
| BPRNACanProMet | [21] |
| BPRNACan3DProMet | [21] |
| CBSX-NSCLC-PBMCs-scRNAseq | [21] |
| CBSX-Melanoma-scRNAseq | [21] |
| CBSX-HNSCC-scRNAseq | [21] |

**Table S3.** Cell types nomenclature for *multideconv*.

| Cell types | Nomenclature |
|---|---|
| B cells | ***B.cells*** |
| B naive cells | ***B.naive*** |
| B memory cells | ***B.memory*** |
| Macrophages | ***Macrophages.cells*** |
| Macrophages type 0 | ***Macrophages.M0*** |
| Macrophages type 1 | ***Macrophages.M1*** |
| Macrophages type 2 | ***Macrophages.M2*** |
| Monocytes | ***Monocytes*** |
| Neutrophils | ***Neutrophils*** |
| Natural Killer cells | ***NK.cells*** |
| Natural killer activated | ***NK.activated*** |
| Natural killer resting | ***NK.resting*** |
| Natural killer T cell | ***NKT.cells*** |
| CD4 T cells | ***CD4.cells*** |
| CD4 naive | ***CD4.naive*** |
| CD4 memory activated | ***CD4.memory.activated*** |
| CD4 memory resting | ***CD4.memory.resting*** |
| CD8 T cells | ***CD8.cells*** |
| Regulatory T cells | ***T.cells.regulatory*** |
| Non regulatory T cells | ***T.cells.non.regulatory*** |

| | |
|---|---|
| Helper T cells | ***T.cells.helper*** |
| Gamma delta T cells | ***T.cells.gamma.delta*** |
| Dendritic cells | ***Dendritic.cells*** |
| Dendritic activated cells | ***Dendritic.activated*** |
| Dendritic resting cells | ***Dendritic.resting*** |
| Cancer cells | ***Cancer*** |
| Endothelial cells | ***Endothelial*** |
| Eosinophils cells | ***Eosinophils*** |
| Plasma cells | ***Plasma*** |
| Myocytes | ***Myocytes*** |
| Fibroblasts | ***Fibroblasts*** |
| Mast cells | ***Mast.cells*** |
| Mast activated cells | ***Mast.activated*** |
| Mast resting cells | ***Mast.resting*** |
| CAF | ***CAF*** |

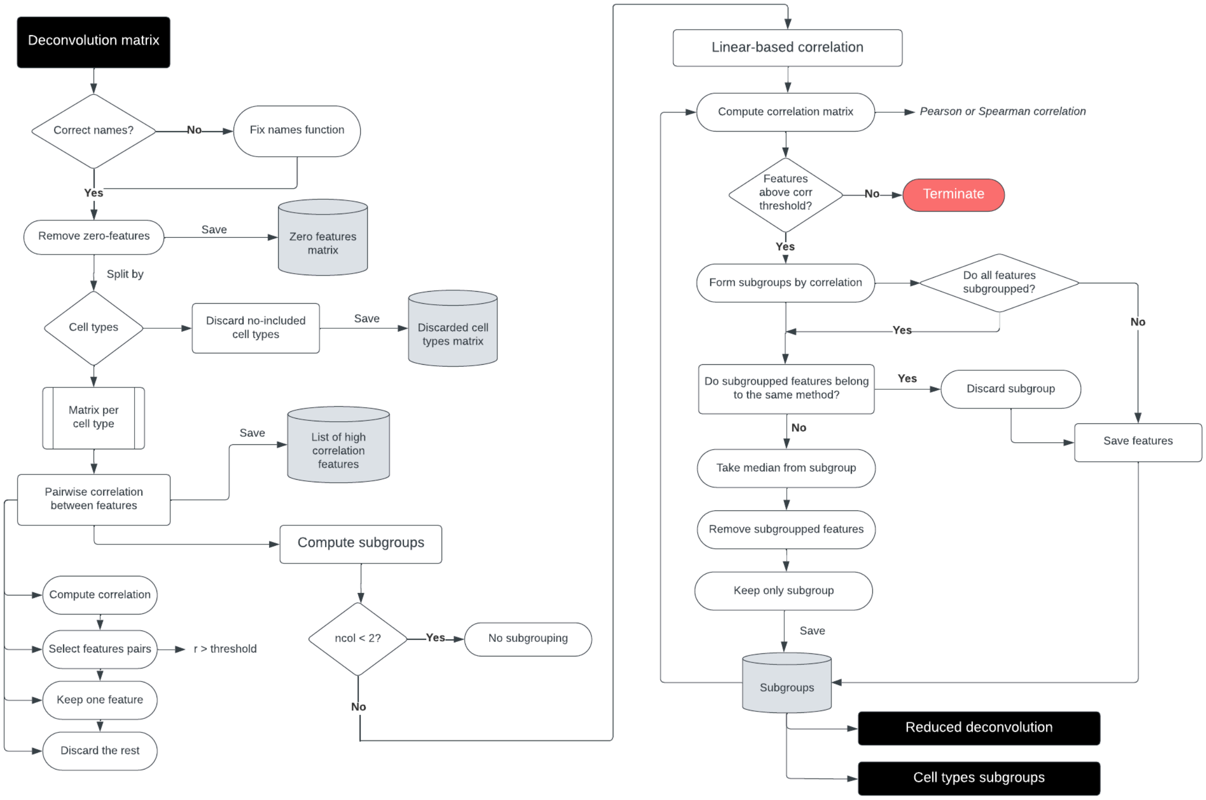


**Figure S1.** Cell-type processing algorithm.

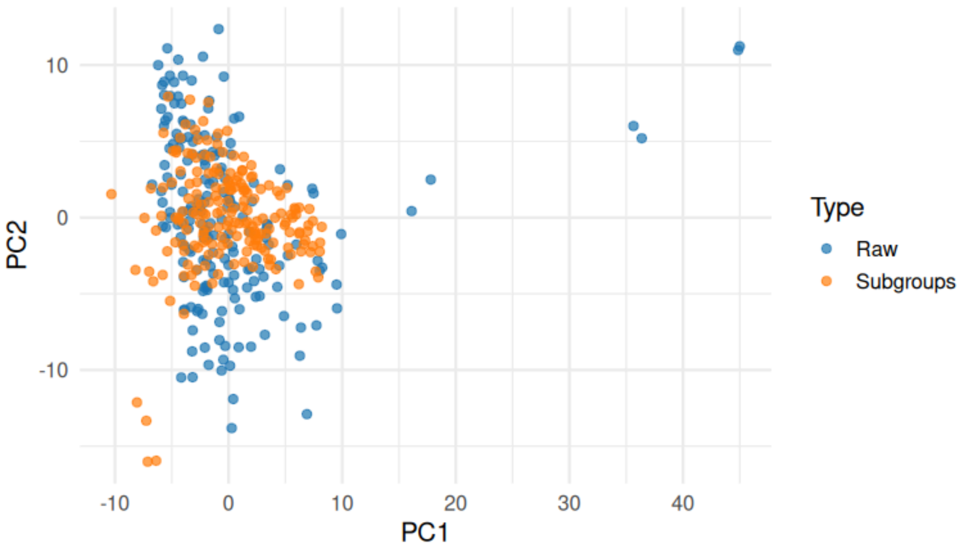


**Figure S2.** PCA using raw deconvolution matrix and subgrouped deconvolution matrix.

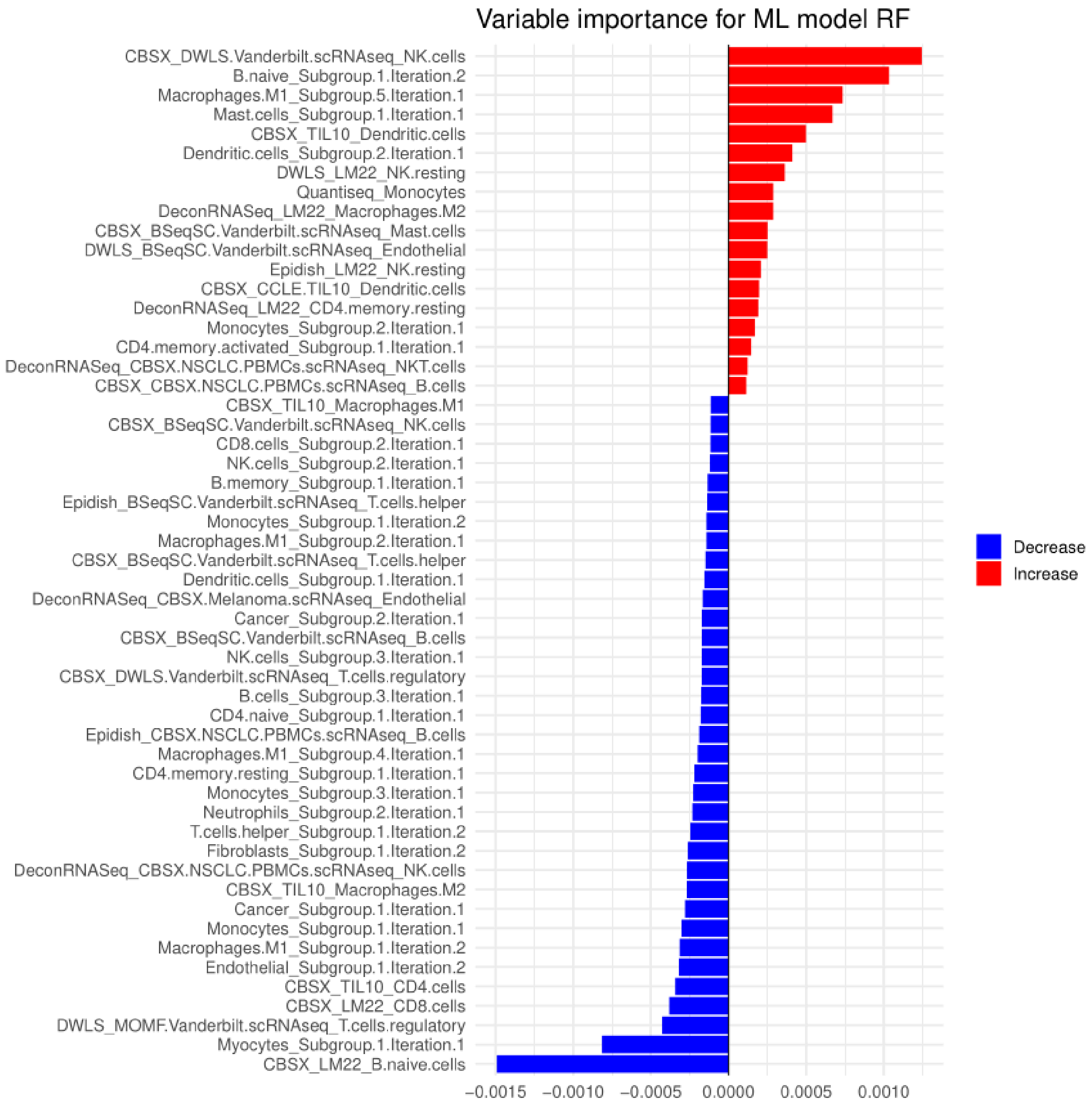


**Figure S3.** Feature importance ranking for the prediction of immunotherapy response using the subgrouped deconvolution features.

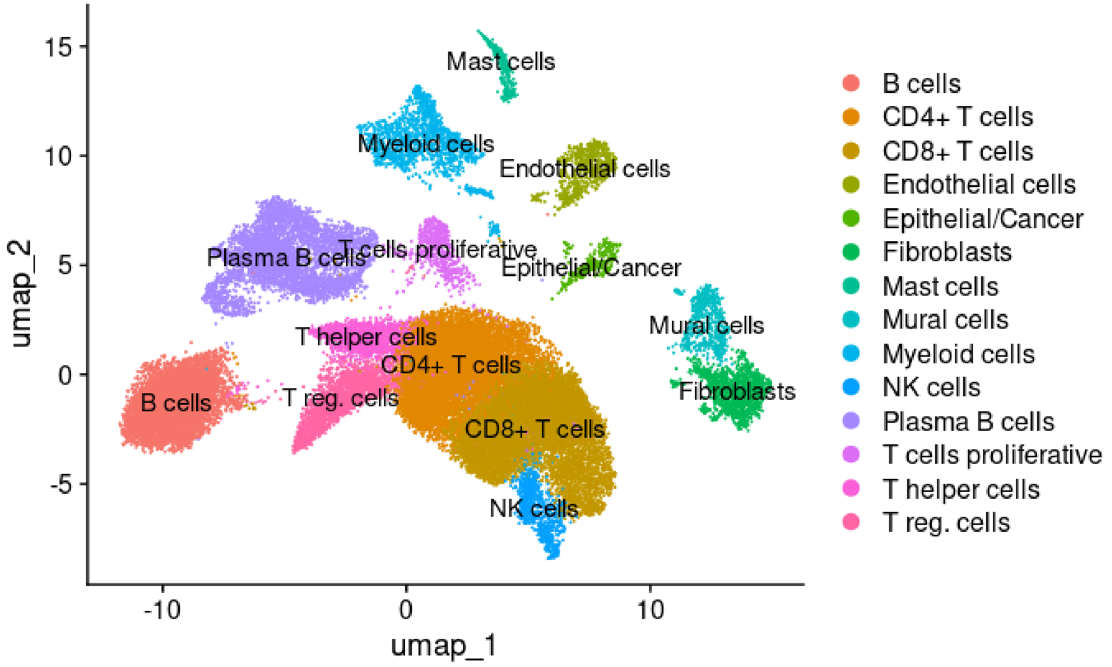


**Figure S4.** UMAP of original scRNA object (Senosain et al. 2023)

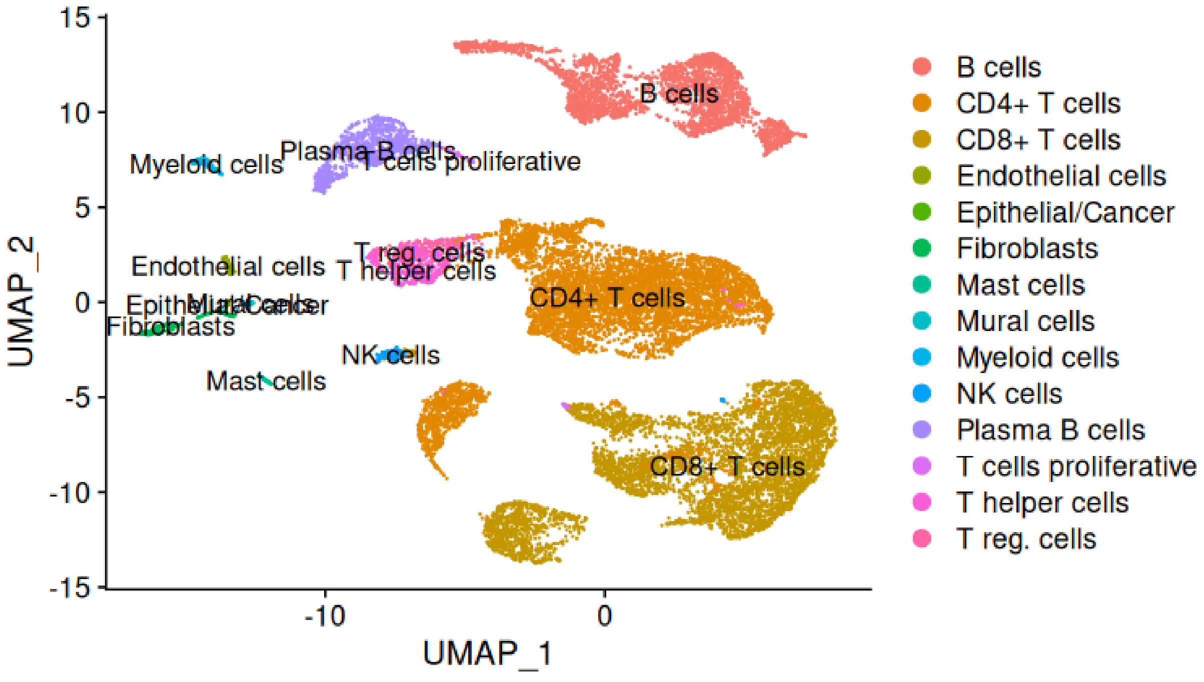


**Figure S5.** UMAP of scRNA object (Senosain et al. 2023) after metacells creation

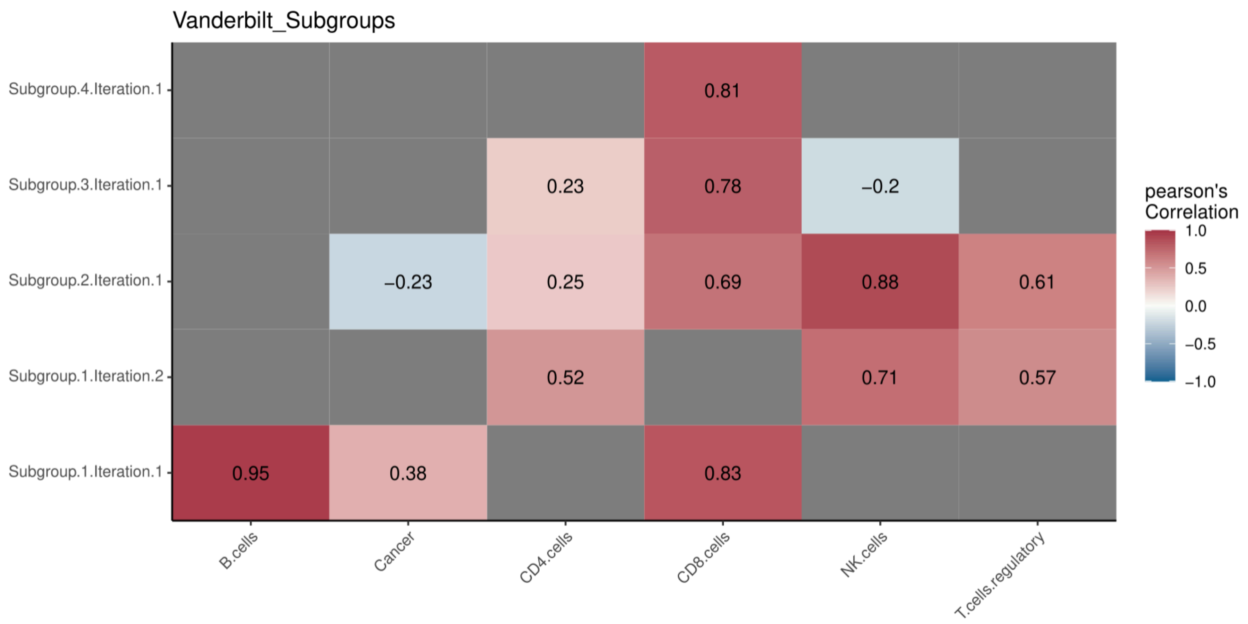


**Figure S6.** Performance of different cell subgroups estimated by *multideconv* (Y-axis) against real cell proportions estimates from the scRNA object on the X-axis (Senosain et al. 2023). Pearson's correlations are shown as colour from -1 (blue) to +1 (red) only when p val<0.05. Non-significant correlations are left unlabeled. Grey boxes indicate cell types that were not estimated because the corresponding signature does not include that cell type.
